# Polaratio: A magnitude-contingent monotonic correlation metric and its improvements to scRNA-seq clustering

**DOI:** 10.1101/2020.12.20.423308

**Authors:** Victor Wang, Pietro Antonio Cicalese, Anto Sam Crosslee Louis Sam Titus, Chandra Mohan

## Abstract

**Motivation:** Single-cell RNA sequencing (scRNA-seq) technologies and analysis tools have allowed researchers to achieve remarkably detailed understandings of the roles and relationships between cells and genes. However, conventional distance metrics, such as Euclidean, Pearson, and Spearman distances, fail to simultaneously take into account the high dimensionality, monotonicity, and magnitude of gene expression data. To address several shortcomings in these commonly used metrics, we present a magnitude-contingent monotonic correlation metric called Polaratio which is designed to enhance the quality of scRNA-seq data analysis.

**Results:** We integrate three interpretable clustering algorithms – Single-Cell Consensus Clustering (SC3), Hierarchical Clustering (HC), and K-Medoids (KM) – through a consensus cell clustering procedure, which we evaluate on various biological datasets to benchmark Polaratio against several well-known metrics. Our results demonstrate Polaratio’s ability to improve the accuracy of cell clustering on 5 out of 7 publicly available datasets.

**Availability:** https://github.com/dubai03nsr/Polaratio

## 1 INTRODUCTION

Single-cell RNA sequencing (scRNA-seq) is a high-throughput transcriptomics technique that allows for the rapid analysis of individual cells in the context of their particular micro-environments (Ziegenhain *et al.,* 2017). This technology has allowed researchers to create a more profound understanding of analyzed tissue samples, ranging from different cell populations, age groups, disease markers, and disease severity (Potter, 2018). The implications for this technique are vast; for example, Bossel Ben-Moshe et *al.* (2019) showed that scRNA-seq could be used to accurately predict tuberculosis in a clinical setting. In addition to diagnostic innovations, scRNA-seq has allowed researchers to identify novel cell populations, which has inspired notable projects such as the Human Cell Atlas (Rozenblatt-Rosen *et al.*, 2017). These advancements were made possible by the development of various unsupervised computational methods, which attempt to cluster the input cells into subgroups based on the relationships between their gene expression values (Wolf *et al.*, 2018; Kiselev *et al.*, 2017; Butler *et al.,* 2018; Becht *et al.,* 2019). In an effort to effectively quantify the relationships between cells, various distance metrics have been explored, including Euclidean, Pearson, and Spearman distances, which can be used to produce a distance matrix for subsequent cell clustering (Kiselev et al., 2019, 2017).

While various techniques have been developed to cope with technical noise and improve data interpretability (Brennecke *et al.*, 2013; McCarthy *et al.,* 2017; Wu and Zhang, 2020), there is still a pressing need for more robust quantitative measures that are able to capture the complex and often non-linear relationships between genes. For example, the expression of genes can be controlled by several different transcription factors which may either work synergistically or antagonistically, while the final transcript levels have varying stabilities and turnover mechanisms. A linear model would be severely handicapped when describing these complex relationships. In addition, the high data dimensionality observed in scRNA-seq data would tend to blur the contrast between small and large inter-cell Euclidean distances (Aggarwal *et al.*, 2001). These data characteristics motivate us to quantify the monotonicity (i.e. the tendency for one variable to increase with another, not necessarily linearly) of the relationship between the expression values in a pair of cells. While the Spearman and Kendall correlations operate on the relative ranks of values in an effort to quantify monotonicity (Hauke and Kossowski, 2010), one shortcoming shared by all rank-based correlations is that in the process of compressing values into their ranks, information derivable from the relative magnitudes of the input values is neglected. To address this shortcoming, we introduce Polaratio, a monotonicity metric that is able to account for the magnitudes composing non-linearities.

## 2 METHODS

Polaratio is designed to yield a high correlation coefficient when the relationship between two inputs follows an arbitrary monotonic relationship (e.g. exponential, logarithmic, etc.), which allows it to effectively represent the complex interactions we observe in gene expression data, such as that shown in Figure 1. While Polaratio’s behavior may resemble that of rank-based correlation metrics, we were interested in also accounting for the *distance* between any two points, which we believe could yield a more robust correlation metric. To accomplish this, we set out to measure the relationship between the sum of the negative and positive monotonic products of the input features for any two cells. Whereas the Spearman correlation coefficient makes use of the differences between ranks, our metric can sensibly describe the magnitude of these differences. To understand why Polaratio fits well in scRNA-seq analysis, we must first consider the nature of the relationship between various discriminative genes and their corresponding cell sub-populations. During proliferation and differentiation of cells, the gradient expression of regulatory genes activates signaling pathways. These gradients are the key markers for identification of cells during these phases, and the magnitude of expression in these genes is important for accurate classification (Bozinovic *et al.,* 2011; Hasegawa *et al.,* 2015). In addition to accounting for this key aspect of scRNA-seq data, Polaratio is scale-invariant (i.e. unaffected by linear scaling) and non-parametric, both of which make Polaratio a versatile distance metric. We provide a graphical abstract of Polaratio and our cell clustering pipeline in Figure 2.

**Fig. 1:**
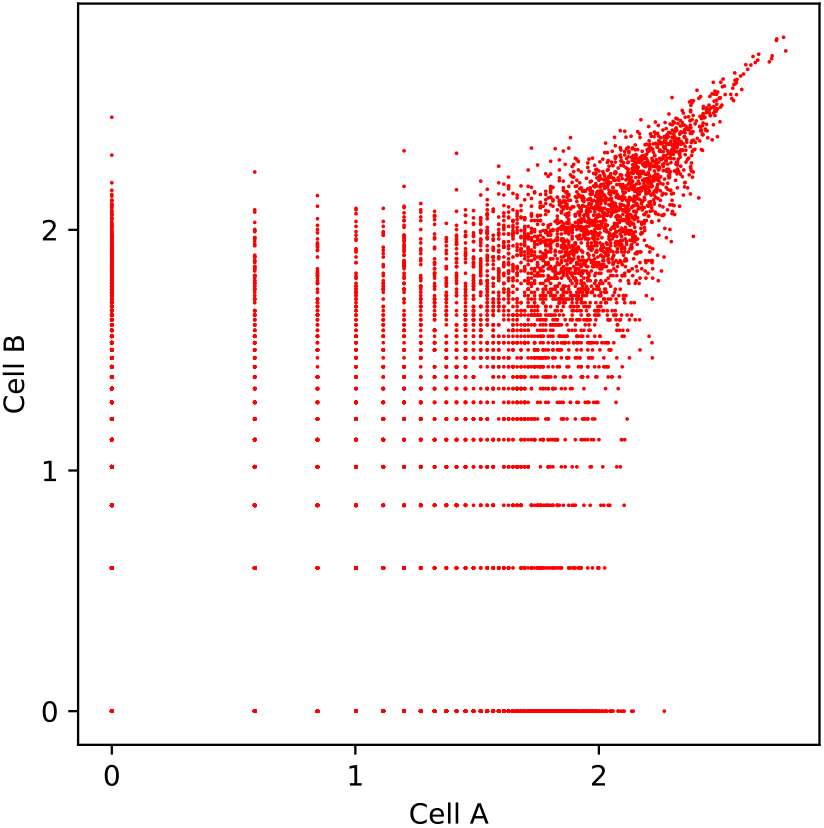
This scatter plot graphs the pre-processed gene expression values for two cells in the two-cell development stage taken from the Goolam dataset (Goolam *et al.,* 2016). The varying concentrations of points along the monotonic trend illustrate that expression magnitude is indeed a noteworthy consideration to incorporate into a distance metric intended to be well-suited for scRNA-seq data. The Polaratio coefficient, as defined in the Methods section, is +0.83 (raw distance = 0.093), the Pearson coefficient is +0.64, and the Spearman coefficient is +0.65.

**Fig. 2:**
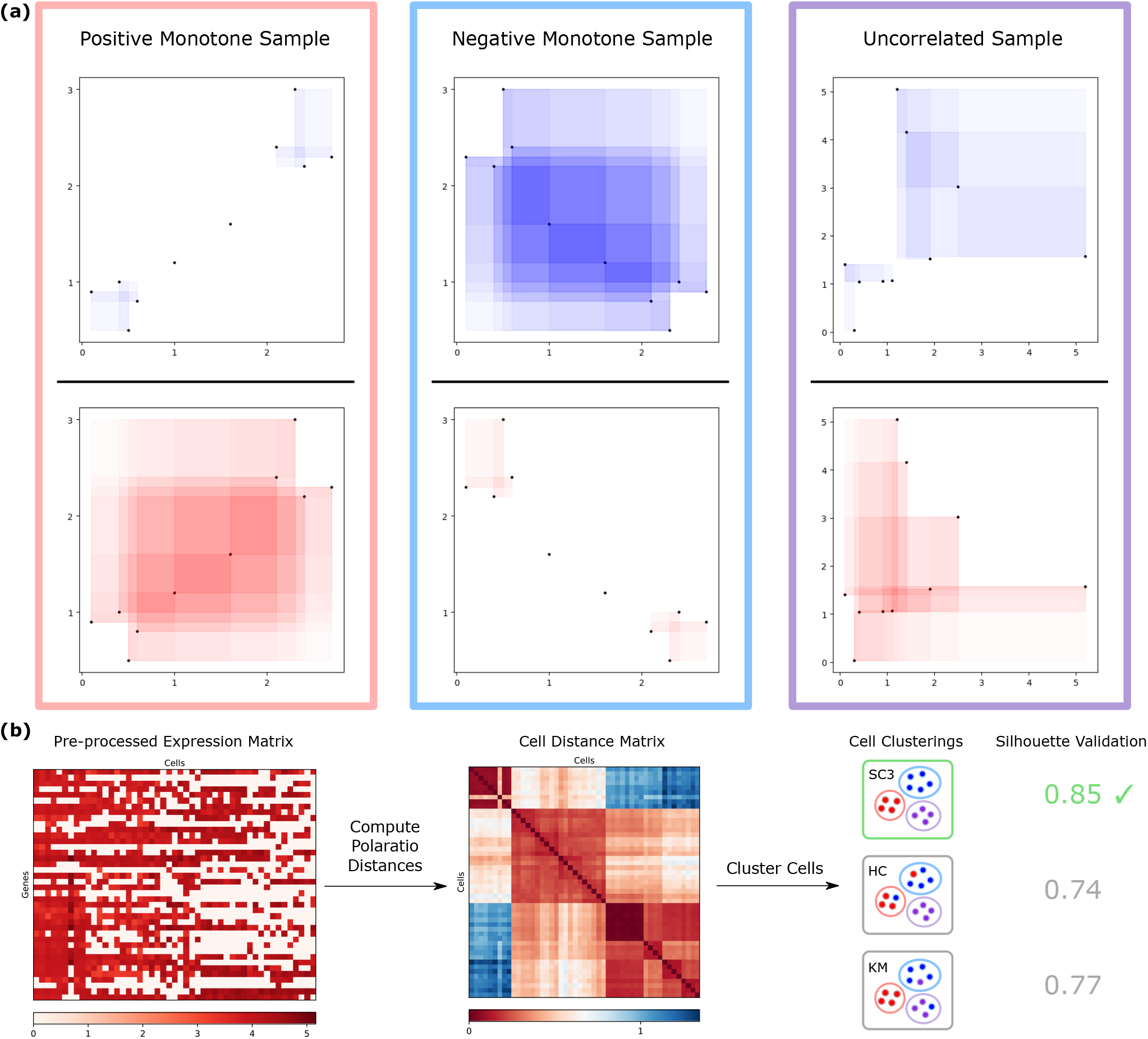
(a): The Polaratio distance between two variables is defined as the ratio of the total area between pairs of points with a negative slope to the total area between pairs of points with a positive slope. Intuitively, the simulation data samples are, from left to right, strongly positively monotonic, strongly negatively monotonic, and nearly uncorrelated. The respective Polaratio coefficients are +0.98, −0.98, and +0.31. In comparison, the corresponding Spearman coefficients are +0.81, −0.81, and +0.67. These sample data illustrate the advantage held by Polaratio over the Spearman correlation: the ability to consider the relations between points in proportion to the magnitudes of their separation. In the positive and negative monotone samples, the relatively minor gaps among the points in the corners are over-magnified by the Spearman correlation’s rank comparisons. Similarly, in the uncorrelated sample, Spearman’s correlation fails to reflect the prominent magnitude of the outliers. **(b)**: To apply Polaratio to cell clustering, we compute the Polaratio distance between every pair of cells with respect to their gene expression values and pass the resulting distance matrix into various clustering algorithms. We then select the top performing clustering algorithm according to the greatest Silhouette index. Note: The raw distances, rather than the correlation coefficients, were used for cell clustering.

### 2.1 Polaratio Distance

We define the Polaratio Distance (PD) as

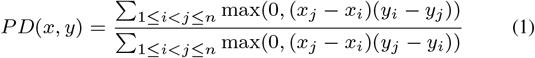

where *x* and *y* represent two variables with *n* observations each. As naively computing the distance would require *O(n^2^)* time, here we outline how we reduced the runtime to *O*(*n* log *n*), matching the time complexity for computing the Spearman correlation, albeit with a higher constant factor. We begin by sorting the points by their *x* values. For a given *j,* efficiently computing

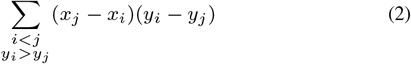

would enable us to quickly compute the numerator. Given that our computations are centered around *j*, we can thus distribute the product (while implying the same summation condition), to produce

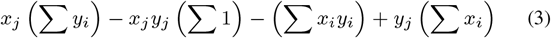

The Binary Indexed Tree (BIT) is a linear-memory data structure that offers two functions, each executed in logarithmic time: (a) to change every value in a specified prefix of an array by a specified amount and (b) to query the sum of the values in a specified prefix of the array (Fenwick, 1994). We update and query a BIT for each of the 4 summations as we sweep through the n points from left to right, resulting in a runtime of *O*(*n* log *n*). The denominator can be similarly expanded and computed, but we can reduce the constant runtime factor by observing that the denominator minus the numerator must equal *n* · COVARIANCE(*x, y*), which can be computed in linear time (see Algorithm 1).

**Algorithm 1.**
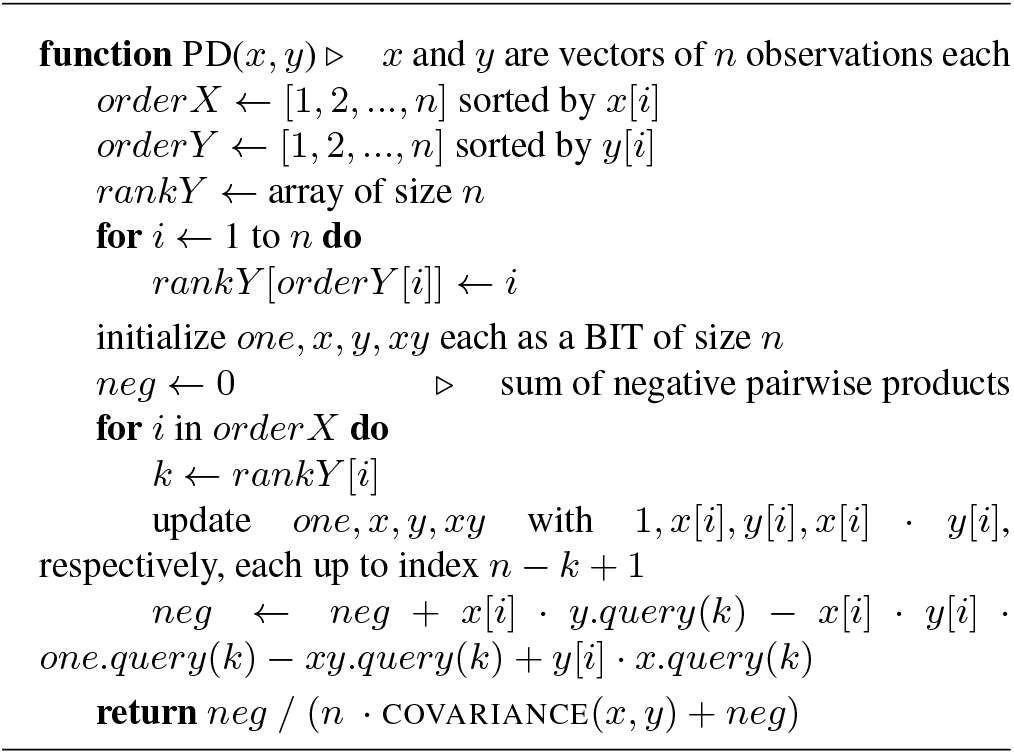
Compute the Polaratio distance between 2 variables

### 2.2 Polaratio Coefficient

An interpretable correlation coefficient is useful for statistical analysis as we may seek to intuitively grasp the relationship between two input samples. To this end, we have developed a function that transforms the Polaratio distance metric into a correlation coefficient within the range [–1,1]. We define the Polaratio Coefficient (PC) following

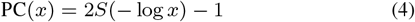

where *x* is the previously defined Polaratio distance, and *S* is the sigmoid function 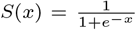 Assuming the natural logarithm, the expression simplifies to

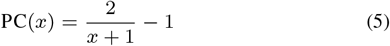

Recall that *x* is the ratio between the total negative products and the total positive products. As such, two relationships that are reflections of each other would yield reciprocal distances and thus coefficients equal in magnitude but opposite in sign, such as the positive and negative monotone samples in Figure 2. In other words, every Polaratio distance *x* satisfies

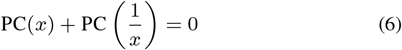

The bounds of the Polaratio distance correspond to the customary bounds of a correlation coefficient: A distance of 0 yields a coefficient of 1 (indicating a perfectly monotone increasing relationship), a distance of 1 yields a coefficient of 0 (indicating a random relationship), and a distance of ∞ yields a coefficient of −1 (indicating a perfectly monotone decreasing relationship).

### 2.3 Clustering Algorithms

The three clustering algorithms we employ in our consensus approach are Single-Cell Consensus Clustering (SC3), complete-linkage agglomerative Hierarchical Clustering (HC), and K-Medoids (KM). We ran SC3 with the R SC3 package, HC with the hclust R method, and KM with the pam method from the R cluster package (Kiselev et al., 2017; R Core Team, 2020; Maechler *et al.,* 2019). SC3 computes distance matrices using the Euclidean, Pearson, and Spearman metrics. The distance matrices are then transformed with principal component analysis and the Laplacian matrix. K-means is then run 1,000 times on varying quantities of retained eigenvectors from the prior transformations. The k-means results are used to generate a consensus matrix, which is then clustered with hierarchical clustering. HC is another frequently used clustering algorithm that allows for intuitive interpretation and offers a flexible choice of number of clusters (Reynolds et al., 2006). The algorithm iteratively combines the two most similar clusters based on a specified agglomeration method (complete-linkage in our approach). Finally, KM has been considered a more robust alternative to k-means by allowing cluster centroids to be represented by certain input objects (Reynolds *et al.*, 2006). These algorithms were appealing options for exploring the limits of Polaratio due to their ability to operate on a specified distance matrix.

### 2.4 Adjusted Rand Index (ARI)

We used the ARI to evaluate how closely each resulting cell clustering matched the one provided by the publishers of the datasets. To calculate the ARI between two clusterings of *n* objects, an *r × s* contingency table *N* is constructed (where the first and second clusterings generated *r* and *s* clusters, respectively) in which *N_i,j_* is the number of objects clustered into cluster *i* and *j* by the first and second clustering, respectively. Letting *a_i_* denote Σ*_J_ N_i,j_* and *b_j_* denote Σ*_J_ N_i,j_*, the ARI is given by

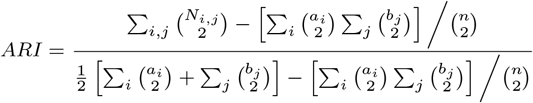

as computed by the adjustedRandIndex method from the R mclust package (Scrucca *et al.*, 2016). A higher ARI indicates a closer match, with an ARI of 1 indicating exact correspondence, and an ARI of 0 indicating random correspondence. Since the number of clusters *k* was specified in every benchmarking trial, r = s was always satisfied.

### 2.5 Silhouette Cluster Validation Index

To optimize performance by determining the “best” clustering out of the three clusterings generated by SC3, HC, and KM, we used the widely accepted Silhouette internal cluster validation index, which relates a given clustering’s inter-cluster separation to its intra-cluster compactness based on the specified distance matrix (Rousseeuw, 1987). The Silhouette index for a given clustering and distance matrix is defined as follows. For a given object *i*, let *a_i_* be the average distance between object *i* and every other object in object *i*’s cluster, and let *b_i_* = min_*C*_ *d*(*i, C*), where *d*(*i, C*) is the average distance between object *i* and every object in a particular cluster *C* to which object *i* does not belong. The Silhouette index *S_i_* for object *i* is then defined as

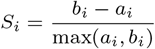

A higher silhouette index indicates a better-clustered object. We define the Silhouette index for a clustering as the average Silhouette index over every object, computed using the silhouette method from the R cluster package (Maechler *et al.*, 2019).

### 2.6 Data Curation

To evaluate the practical significance of Polaratio, we chose datasets of various tissue domains, dataset sizes, numbers of sub-populations, and sampling units. Our dataset selection process follows what was proposed in (Kiselev et al., 2017), where a gold and silver standard was adopted to appropriately consider the results in light of the objectivity of the cell labels. We consider the Biase, Goolam, Yan, Deng, Pollen, and Bozec datasets to be gold standards because the cell sub-populations were chosen from distinct biological stages or conditions (Biase *et al.,* 2014; Goolam *et al.,* 2016; Yan et al., 2013; Deng et al., 2014; Pollen et al., 2014; Andreev et al., 2020). We consider the Treutlein dataset to be a silver standard because the cells were labeled according to the authors’ computational analysis and biological knowledge (Treutlein *et al.,* 2014). Table 1 provides a detailed list of each dataset’s attributes.

**Table 1.** A summary of the datasets. *k* is the number of cell clusters, n is the number of cells analyzed, and *m* is the number of genes after pre-processing.

| Dataset | $k$ | $n$ | $m$ | Units |
| --- | --- | --- | --- | --- |
| Biase | 3 | 49 | 6323 | FPKM |
| Goolam | 5 | 124 | 11157 | CPM |
| Yan | 8 | 124 | 9460 | RPKM |
| Deng | 10 | 259 | 10372 | RPKM |
| Pollen | 11 | 301 | 11828 | TPM |
| Bozec | 2 | 1016 | 6317 | CPM |
| Treutlein | 4 | 201 | 9462 | FPKM |

## 3 RESULTS AND DISCUSSION

### 3.1 Pre-processing

We filtered out rare and ubiquitous genes (genes not expressed in more than 90% or less than 10% of the cells) with the methods `sc3_prepare` and `get_processed_dataset` from the SC3 R package, due to the lack of discriminative information derived from such genes (Kiselev et al., 2017). The filtered matrix was then log-transformed and normalized using the logNormCounts method from the scater R package to further mitigate the impact of noise on our analysis (Amezquita et al., 2020; McCarthy et al., 2017). To prevent the artificial improvement of our results due to genes recorded exclusively from distinct cell populations, we only considered the genes whose expression values were recorded across all cell sub-populations. This condition was not required for the Yan dataset, in which all of the genes whose expression values were recorded across all cells were ubiquitous genes (presumably because expression values of 0 were preemptively removed).

### 3.2 SC3 with Individual Metrics

We chose to benchmark our metric with SC3 because it has been shown to be among the most robust state-of-the-art cell clustering pipelines for scRNA-seq data (Duó A and C., 2020). We ran SC3 on the above datasets to compare Polaratio to the Euclidean, Pearson, and Spearman distances, which SC3 uses by default. To compare the individual metrics, we ran SC3 with each metric alone, as opposed to SC3’s default consensus setting. We did this to gain an understanding of the instances where Polaratio could contribute a tangible benefit to our clustering analysis. Although SC3 is relatively stable compared to other cell clustering algorithms, its performance may still experience fluctuation from run to run, so we repeated each trial 10 times and recorded the average performance. Table 2 lists the performance of the individual metrics through SC3, with green text indicating the statistically significant highest mean ARI (Wilcoxon signed-rank test p-value < 0.01). We note that SC3 with Polaratio achieves statistically superior results in the Biase, Deng, and Bozec datasets.

**Table 2.** SC3 Mean ARI with Individual Metrics

| Dataset | Polaratio | Euclidean | Pearson | Spearman |
| --- | --- | --- | --- | --- |
| Biase | 1.000 | 0.948 | 0.948 | 0.948 |
| Goolam | 0.427 | 0.687 | 0.687 | 0.687 |
| Yan | 0.795 | 0.786 | 0.795 | 0.770 |
| Deng | 0.655 | 0.481 | 0.576 | 0.611 |
| Pollen | 0.952 | 0.962 | 0.960 | 0.979 |
| Bozec | 0.568 | 0.307 | 0.475 | 0.559 |
| Treutlein | 0.640 | 0.606 | 0.637 | 0.840 |

### 3.3 Comparison of Consensus Approaches

As Polaratio is a generic metric in that it can be applied broadly, we explored its usage in two more clustering algorithms, HC and KM. In doing so, we observed that each clustering algorithm performed well on some datasets but poorly on others, so we used the Silhouette index to determine the “best” clustering. In each trial, we selected the clustering with the highest Silhouette index as our final clustering. HC and KM were run exactly once per trial because they are deterministic, but due to SC3’s stochasticity, we ran SC3 5 times per trial and retained the clustering with the highest Silhouette index. We reduced the number of runs from 10 in the previous section to 5 in this section because as only the “best” instead of the mean performance was analyzed, we opted to minimize the unnecessary computational cost. The above consensus approach was performed on each metric individually to allow for fair comparison between them. We observed superior performance on 5 out of the 7 datasets with the Polaratio consensus approach, and only a marginal decrease in the Pollen dataset (see Table 3). Therefore, we conclude that Polaratio is more successful than popular metrics at distinguishing cell labels with gene expression data.

**Table 3.** Comparison of default SC3 and each metric’s performance in the consensus approach. Bold values indicate the ARI of the final clustering selected by the highest Silhouette index. The final column offers the default SC3 pipeline (which incorporates the three metrics Euclidean, Pearson, and Spearman) as a baseline benchmark, in which each value represents the mean ARI over 10 trials. Although our consensus approach is nearly deterministic, in cases where the SC3 clustering was selected, we introduced the standard deviation of the ARI from the 10 SC3 trials with the corresponding metric(s) to allow for statistically fair comparison. Green values indicate a statistically significant higher mean ARI (Wilcoxon signed-rank test p-value < 0.01) compared to the default SC3 as well as the other selected clusterings (note that no green text appears for the Deng dataset because the Pearson SC3 performance was not observed to be statistically superior to the default SC3 performance).

| Dataset | Polaratio |  |  | Euclidean |  |  | Pearson |  |  | Spearman |  |  | SC3 |
| --- | --- | --- | --- | --- | --- | --- | --- | --- | --- | --- | --- | --- | --- |
|  | HC | KM | SC3 | HC | KM | SC3 | HC | KM | SC3 | HC | KM | SC3 |  |
| Biase | **1.000** | **1.000** | **1.000** | 1.000 | 1.000 | **0.948** | 1.000 | 1.000 | **0.948** | 1.000 | 1.000 | **0.948** | 0.948 |
| Goolam | **0.914** | 0.894 | 0.427 | **0.711** | 0.732 | 0.687 | **0.656** | 0.738 | 0.687 | **0.595** | 0.761 | 0.687 | 0.687 |
| Yan | 0.742 | **0.811** | 0.795 | **0.672** | 0.811 | 0.786 | **0.642** | 0.811 | 0.795 | 0.703 | 0.719 | **0.770** | 0.795 |
| Deng | **0.459** | 0.350 | 0.583 | **0.382** | 0.411 | 0.482 | 0.393 | 0.414 | **0.559** | **0.366** | 0.433 | 0.608 | 0.556 |
| Pollen | 0.558 | 0.811 | **0.953** | 0.541 | 0.793 | **0.960** | 0.286 | 0.811 | **0.960** | 0.586 | 0.829 | **0.977** | 0.961 |
| Bozec | 0.442 | 0.533 | **0.571** | 0.009 | 0.530 | **0.213** | 0.556 | 0.521 | **0.469** | 0.559 | **0.504** | 0.559 | 0.530 |
| Treutlein | **0.812** | 0.449 | 0.640 | 0.674 | **0.103** | 0.606 | **0.777** | 0.547 | 0.637 | **0.330** | 0.544 | 0.840 | 0.647 |

## 4 CONCLUSION

We have presented a novel distance metric, Polaratio, that expresses the monotonic tendency between two variables. Polaratio consolidates many important characteristics of common distance metrics, rendering it capable of capturing non-linearities while factoring in the magnitude of the distances between observations. Our benchmarking showed that Polaratio outperforms popular distance metrics at cell clustering when we combine multiple clustering algorithms with our Silhouette index selection criterion. Future works should explore methods that assign weights to genes as a means to further improve clustering quality. Additionally, further analysis should be conducted to make better sense of the underlying data properties that allow Polaratio to succeed. Finally, as Polaratio does not rely on any properties exclusive to scRNA-seq data, its use should be investigated in other domains to improve general statistical analysis.

## 5 DATA AVAILABILITY

Biase: ncbi.nlm.nih.gov/geo/query/acc.cgi?acc=GSE57249 – The 3 cell labels used were zygote, 2-cell, and 4-cell, for a total of 49 cells.

Goolam: ebi.ac.uk/arrayexpress/experiments/E-MTAB-3321/ - The 5 cell labels used were 2-cell, 4-cell, 8-cell, 16-cell, and 32cell, for a total of 124 cells.

Yan: ncbi.nlm.nih.gov/geo/query/acc.cgi?acc=GSE36552 - The 8 cell labels used were oocyte, zygote, 2-cell embryo, 4-cell embryo, 8-cell embryo, morulae, late blastocyst, and human embryonic stem cell (hESC), for a total of 124 cells.

Deng: ncbi.nlm.nih.gov/geo/query/acc.cgi?acc=GSE45719 – The 10 cell labels used were zygote, early 2-cell, mid 2-cell, late 2-cell, 4-cell, 8-cell, 16-cell, earlyblast, midblast, and lateblast, for a total of 259 cells.

Pollen: hemberg-lab.github.io/scRNA.seq.datasets/human/tissues/ – The 11 cell labels used were 2338, 2339, K562, BJ, HL60, iPS, Kera, GW21.2, GW21, NPC, and GW16, for a total of 301 cells. Bozec: ncbi.nlm.nih.gov/geo/query/acc.cgi?acc=GSE153632 – The 2 cell labels used were Lung_OVA+SIA and Synovium_OVA+SIA, for a total of 1016 cells.

Treutlein: ncbi.nlm.nih.gov/geo/query/acc.cgi?acc=GSE52583 – The 4 cell labels used were E14.5, E16.5, E18.5, and adult, for a total of 201 cells.

## Notes

https://www.ncbi.nlm.nih.gov/geo/query/acc.cgi?acc=GSE57249

https://www.ebi.ac.uk/arrayexpress/experiments/E-MTAB-3321/

https://www.ncbi.nlm.nih.gov/geo/query/acc.cgi?acc=GSE36552

https://www.ncbi.nlm.nih.gov/geo/query/acc.cgi?acc=GSE45719

https://hemberg-lab.github.io/scRNA.seq.datasets/human/tissues/

https://www.ncbi.nlm.nih.gov/geo/query/acc.cgi?acc=GSE153632

https://www.ncbi.nlm.nih.gov/geo/query/acc.cgi?acc=GSE52583

### Competing Interest Statement

The authors have declared no competing interest.

### Summary of Updates

Correction in Figure 2.

